# Resolving brain organoid heterogeneity by mapping single cell genomic data to a spatial reference

**DOI:** 10.1101/2020.01.06.896282

**Authors:** Jonas Simon Fleck, Zhisong He, Michael James Boyle, J. Gray Camp, Barbara Treutlein

**Affiliations:** Department of Biosystems Science and Engineering, ETH Zürich, Basel, Switzerland; Max Planck Institute for Evolutionary Anthropology, Leipzig, Germany; Institute of Molecular and Clinical Ophthalmology, Basel, Switzerland

## Abstract

Self-organizing tissues resembling brain regions grown in vitro from human stem cells (so-called organoids or spheroids) offer exciting possibilities to study human brain development, disease, and evolution. Brain organoids or spheroids are complex and can contain cells at various stages of differentiation from different brain structures. Single-cell genomic methods provide powerful approaches to explore cell composition, differentiation trajectories, gene regulation, and genetic perturbations in brain organoid systems. However, it remains a major challenge to understand the cellular heterogeneity observed within and between individual organoids. Here, we have developed a computational approach (VoxHunt) to assess brain organoid patterning, developmental state, and cell composition through systematic comparisons to three-dimensional *in situ* hybridization data from the Allen Brain Atlas. Cellular transcriptomes as well as accessible chromatin landscapes can be compared to spatial transcript patterns in the developing mammalian brain, which enables characterization and visualization of organoid cell compositions. VoxHunt will be useful to assess novel organoid engineering protocols and to annotate cell fates that emerge in organoids during genetic and environmental perturbation experiments.

## INTRODUCTION

Spatiotemporal patterning events occurring during brain development lead to the formation of multiple forebrain, midbrain, and hindbrain structures composed of a diversity of cell types. Portions of this structural and cellular diversity can be recapitulated in stem cell-derived neuronal tissues that self-organize in three-dimensional culture *in vitro* (Lancaster and Knoblich, 2014). Over the past few years, the generation of human brain organoids and spheroids has enabled unprecedented analyses of human brain development and disease (reviewed in (Arlotta, 2018; Giandomenico and Lancaster, 2017; Pasca, 2018), as well as evolution (Kanton et al., 2019; Mora-Bermúdez et al., 2016; Pollen et al., 2019). There are two general strategies for producing 3D human brain tissue cultures. In the first strategy, pluripotent stem cells are guided to form a layer of neuroepithelial stem cells that can differentiate into neurons. Culturing the neuroepithelium in permissive media can result in the generation of multiple brain structures (Figure 1a). This self-patterning strategy (Lancaster et al., 2013; Sasai, 2013) offers the potential to understand how brain structures self-organize and interact. However, there are often substantial differences between individual organoids, and between batches grown separately. The alternative strategy is to use signaling molecules to control patterning of the neuroepithelium so that a defined structure forms, such as the cortex (Eiraku et al., 2008; Kadoshima et al., 2013; Pasca et al., 2015), ventral telencephalon (Bagley et al., 2017; Birey et al., 2017; Xiang et al., 2017), hypothalamus (Qian et al., 2016), thalamus (Xiang et al., 2019), and others. Multiple protocols have been published to generate modular brain structures and to create higher-level structures by fusing two or more brain structures to study neuronal migration or to enable formation of inter-region neuronal connections (Bagley et al., 2017; Birey et al., 2017; Pasca, 2018; Xiang et al., 2019). This technique might increase reproducibility, but researchers have yet to define all of the signals that create each brain structure. Moreover, it is unclear whether certain structures can form in the absence of adjacent structures. Despite incredible progress in human brain organoid engineering, it is not yet possible to create a stereotypic organoid containing multiple brain structures.

**Figure 1:**
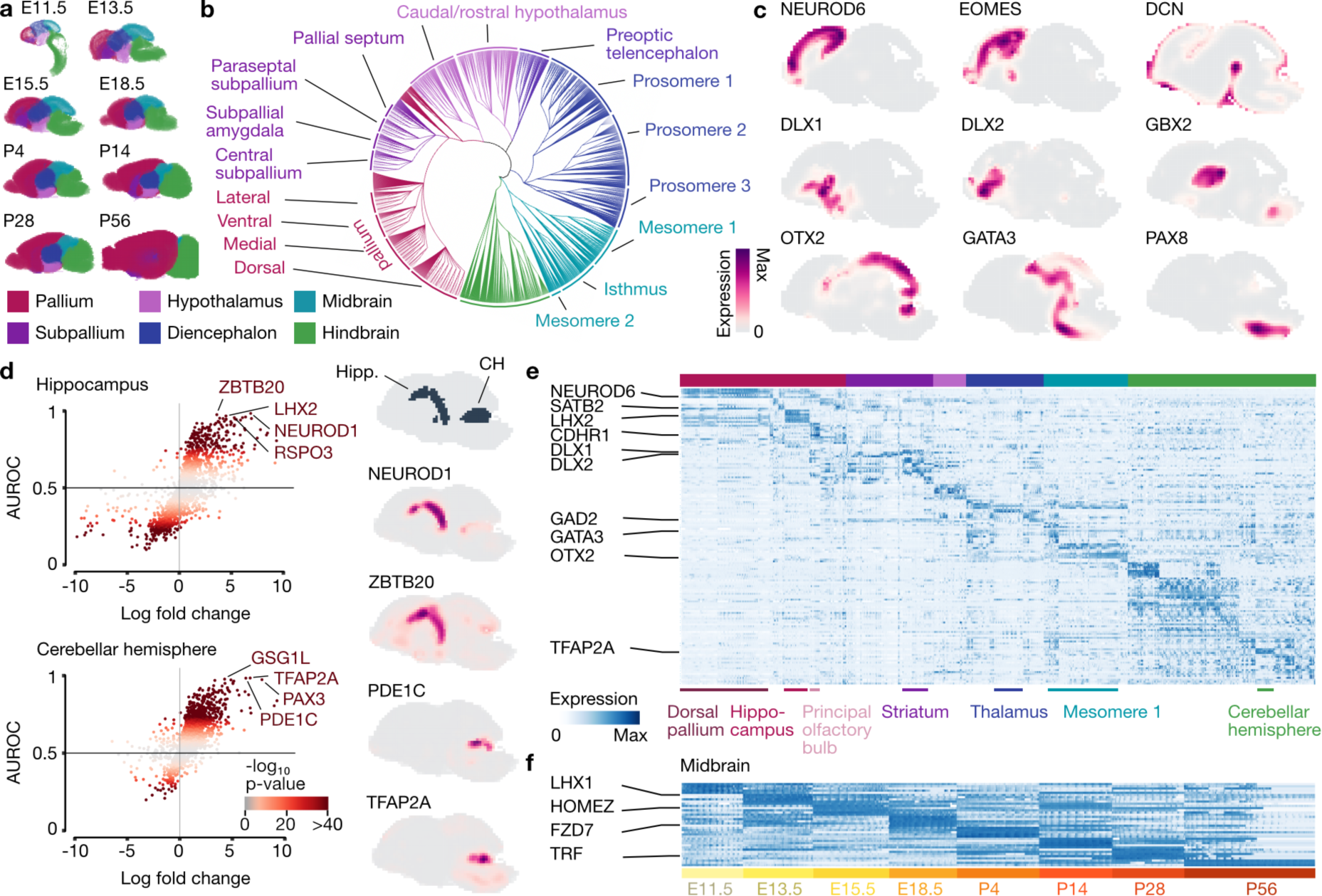
Feature extraction from voxel maps of developing mouse brain in situ hybridization data. a) Schematics show mouse brain development over time, with high-level structure annotations color-coded. b) Dendrogram showing the hierarchically organized structure ontology from the Allen Developing Mouse Brain Atlas. c) Spatial expression patterns of selected markers as maximum intensity projections across five central sagittal sections (8-12) in the E15.5 mouse brain. d) Scatterplot showing feature selection for two brain structures, Cerebellar hemisphere (CH) and Hippocampus (Hipp.) at E15.5, together with the expression profile for the top two marker genes per structure. e-f) Heatmap showing marker gene expression across brain structures at E15.5 (e), and across developmental stages for the midbrain (f).

Organoid protocol development requires robust strategies to assess organoid composition. Single-cell mRNA sequencing (scRNA-seq) has emerged as a high-resolution approach to deconstruct organoid cell composition and reconstruct differentiation trajectories in these complex developing tissues (Camp et al., 2015; Kanton et al., 2019; Quadrato et al., 2017; Velasco et al., 2019). Multiple labs have used scRNA-seq to assess heterogeneity in brain organoids, and direct comparison of cortical structures to primary counterparts have revealed remarkable similarity in gene expression patterns between the *in vitro* and *in vivo* tissues (Camp et al., 2015; Pollen et al., 2019). However, it still remains a challenge to interpret gene expression patterns from organoid single-cell genomic data. Here, we have developed a computational approach to assess brain organoid patterning, developmental state, and cell composition through systematic comparisons to *in situ* hybridization data from the Allen Brain Atlas. Based on their transcriptome or accessible chromatin landscape, cells can be mapped to voxels, in this case 3D positions rendered from spatial transcript patterns, which enables characterization and visualization of organoid cell composition. VoxHunt will be useful to assess novel organoid engineering protocols and to rapidly annotate cell fates that emerge in organoids during genetic and environmental perturbation experiments.

## RESULTS

To facilitate exploration of 3D gene expression patterns in the developing brain, we obtained *in situ* hybridization data of the mouse brain in 8 different developmental stages (Figure 1a) from the Allen Developing Mouse Brain Atlas (Thompson et al., 2014). This dataset provides spatial expression patterns from different experiments that measure a total of 2,073 genes, which are registered to a single voxel grid for each timepoint (80 - 200 µm resolution) (Figure S1). Each voxel is further annotated with a brain structure derived from a hierarchical structure ontology. We preprocessed this data by summarizing *in situ* hybridization quantifications from different experiments measuring the same gene to obtain a single gene expression vector for each voxel at each developmental stage. To allow for comparison of structures with similar size throughout developmental stages, we further introduced additional layers of structure annotation by aggregating existing ontological levels, resulting in spatially and temporally resolved brain structures (Figure 1b-c, Figure S2).

We performed differential expression analysis between annotated brain structures and used the single-feature area under the receiver operating characteristic curve (AUROC) as a metric for selecting structure-specific genes with spatially confined expression (Figure 1d, Figure S3a). In this way, one can perform large-scale feature selection based on various criteria such as structure- or stage-specificity (Figure 1e-f, Figure S3b). These brain structure-enriched gene expression profiles could serve as a resource of marker genes to validate novel cell fate engineering and organoid protocols.

Next, we tested if this 3D spatial *in situ* dataset could be leveraged for unsupervised annotation of cell types generated through different organoid protocols. For this, we obtained published single-cell data from three brain structure-specific organoid protocols (thalamus, cortex, ventral telencephalon) (Birey et al., 2017; Xiang et al., 2019) as well as one cerebral (unpatterned) organoid protocol (Kanton et al., 2019). We used Uniform Manifold Approximation and Projection (UMAP) (Becht et al., 2018) embeddings to project cells onto a two-dimensional space for each protocol, and annotated the clusters as described in the original studies (Figure 2a). All four datasets exhibit a diversity of cell populations, including heterogeneous progenitor and neuronal cells. For neuronal cells from structure-specific organoids, successful patterning is supported by the expression of canonical marker genes (Figure 2b). For cerebral organoids, multiple brain structures can form in any given organoid, and marker gene expression analysis suggests the emergence of neuronal cell types from the telencephalon (pallium, subpallium), diencephalon, midbrain, and hindbrain supporting the annotations reported in the original publication (Figure 2b). We assessed if annotations from each dataset could be confirmed by correlating single cell transcriptomes from each neuronal cluster with *in situ* expression patterns in the mammalian brain at high-level annotations that are consistent across time for telencephalon (pallium, subpallium), diencephalon, midbrain, and hindbrain. Cells annotated as cortical neurons derived from both cortical or cerebral organoid protocols showed far higher correlation to pallium than to any other structure. Thalamic neurons showed a similarly clear pattern, with the majority (>90%) of cells having the highest correlation to voxels within the diencephalon. For clusters of inhibitory neurons annotated as ventral telencephalon, we observed differences between the two organoid protocols. While cells derived from cerebral organoids had consistently higher correlation with subpallial structures, a large fraction of cells (48.3%) from ventrally patterned organoids were most highly correlated with the hypothalamus. Cell clusters annotated as midbrain neurons derived from cerebral organoid protocols showed a noisier correlation pattern and overall lower correlation to *in situ* maps than any of the other clusters. While average correlation within each cluster was highest to the midbrain, correlation of single cells did not show clear agreement, especially for cluster 6 annotated as ‘midbrain excitatory neurons’.

**Figure 2:**
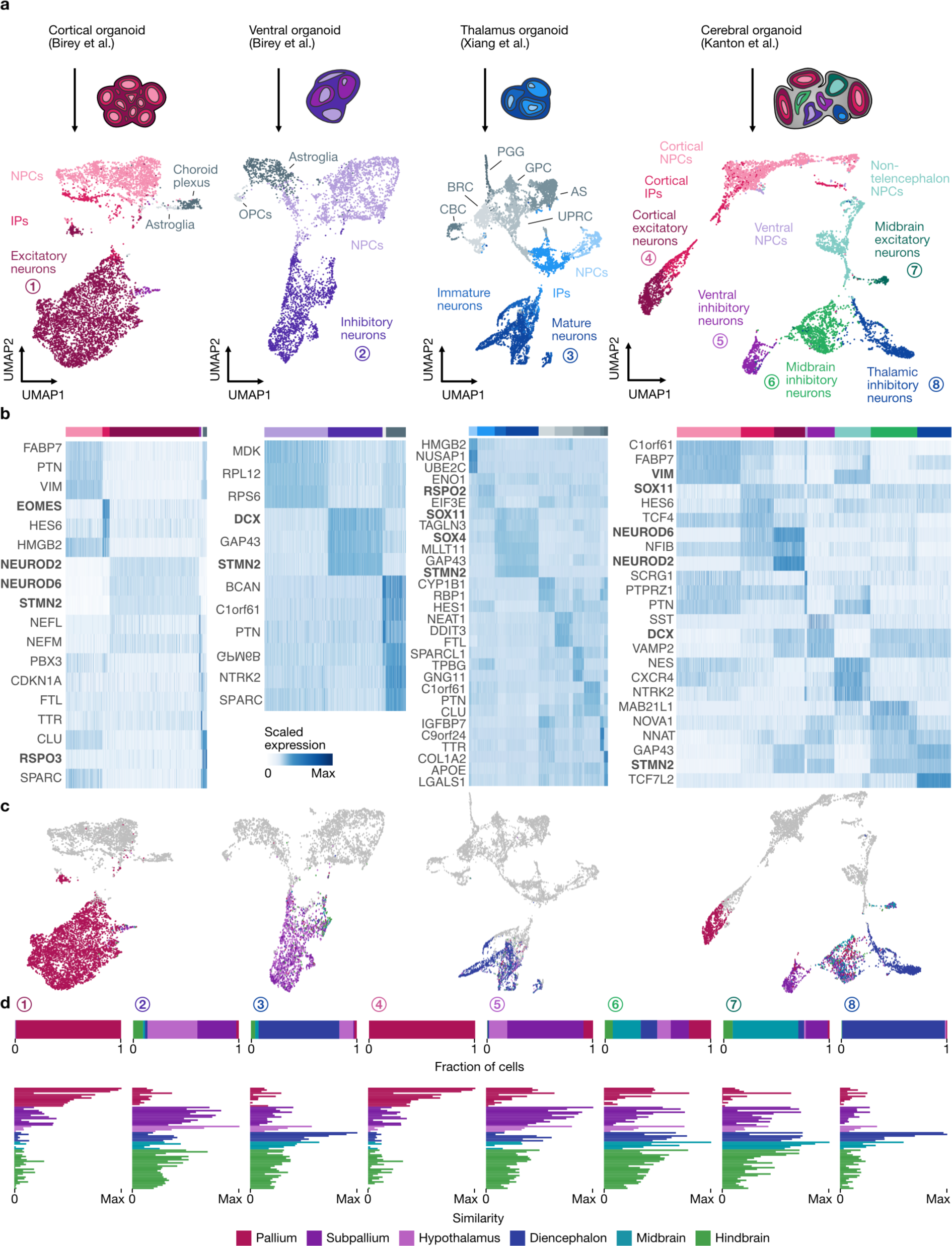
Single-cell transcriptomes deconstruct cell composition in brain organoids. a) UMAP projections of published single-cell transcriptome data from from different organoid protocols (cortical (Birey et al., 2017), ventral (Birey et al., 2017), thalamus (Xiang et al., 2019), cerebral (Kanton et al., 2019)). Cells are colored based on the annotations in the original publications. b) Heatmap showing the expression of top marker genes for each cluster. Canonical marker genes are highlighted in bold. c) UMAP projections of neuronal cells from each dataset colored by structure annotation of the voxel with maximum correlation. d) Top, Fraction of cells from each cluster annotated in panel a that have maximum correlation to each brain structure based on comparison to *in situ* hybridization voxel maps. Bottom, bar plots showing the similarity of each cluster to different brain structures. IP: Immediate progenitor cells, NPC: Neural progenitor cells, OPC: Oligodendrocyte progenitor cells, CBC: Cilia-bearing cells, BRC: BMP-related cells, PGG: Proteoglycan-expressing glia, GPC: Glial progenitor cell, AS: Astrocytes, UPRC: Unfolded protein response-related cells.

To select a suitable developmental stage for detailed voxel-resolved visualization of spatial similarity patterns, we computed the correlation of each cell in a cluster to its matching structure from all fetal developmental stages (E11.5 -E18.5) (Figure 3a, Figure S5). Most clusters showed the highest median correlation to E13.5, with the exception of thalamic neuron clusters, which were most highly correlated with structures at E11.5 (Figure 3b). For each organoid cluster, we visualized the correlation to each voxel in either a sagittal view (Figure 3c, Figure S5) or two-dimensional coronal (Figure 3d) sections. To further aid the assessment of cell type compositions of individual organoids, each cell from each organoid can be additionally projected to spatial locations within the brain based on maximum similarity to loci within the voxel maps (Figure 3e). These analyses reveal the spatial location within the brain tissue where each cluster or cell has maximal correlation, which can be used to assess the specificity of the regional neuronal population and help annotate previously unknown cell populations (Figure S4). Correlation patterns can also be explored in three-dimensional registrations of the voxel maps (Videos S1-9).

**Figure 3:**
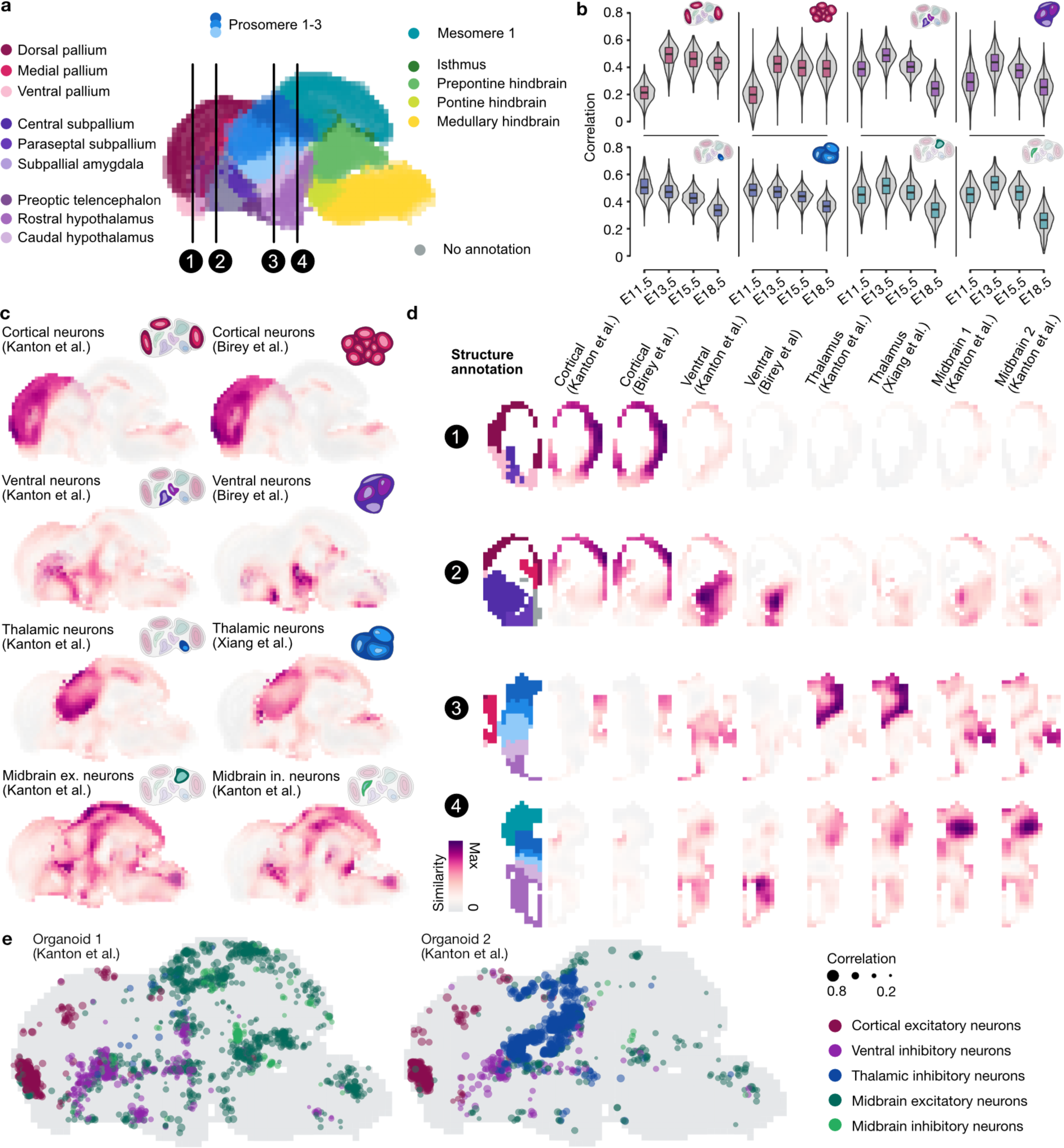
Comparing brain organoid cell populations to spatial brain maps. a) Structure annotations of the E13.5 mouse brain (sagittal view), which was used for organoid cluster annotation. Locations 1-4 are used for coronal sections shown in subsequent panels. b) Violin plots showing the organoid cluster correlation distributions to voxel maps of the respective structure (pallium, subpallium, diencephalon, mesencephalon) across different developmental time points. c) Sagittal projections colored by scaled similarity scores of different organoid clusters from each dataset to voxel maps of the E13.5 mouse brain. d) Coronal sections visualizing regional annotation and scaled similarity scores to different brain regions from different clusters and protocols. e) Sagittal view of a maximum-correlation projection of cells from two individual cerebral organoids to 8 central sections (5-12) of the E13.5 mouse brain.

Finally, we show that similar analyses can be performed for single-cell ATAC-seq (Assay for Transposase-Accessible Chromatin using sequencing) data. In this case, we obtained paired scATAC-seq and scRNA-seq data performed on the same cell suspension generated from microdissected regions from 2-4 month cerebral organoids (Kanton et al., 2019) (Figure 4a). We determined accessibility scores at the transcription start site (TSS) of each gene measured in the *in situ* hybridization atlas (Figure 4b), and then calculated TSS access and expression correlation across each structure annotated in the voxel map (Figure 4c-d). Coloration of the 3D voxel map based on correlation revealed that both transcriptomes and chromatin accessibility scores of neuronal cells from the microdissected region have the highest correlation with the developing cortex (Figure 4c). This demonstrates that VoxHunt can consistently annotate cells across multiple single-cell -omics modalities.

**Figure 4:**
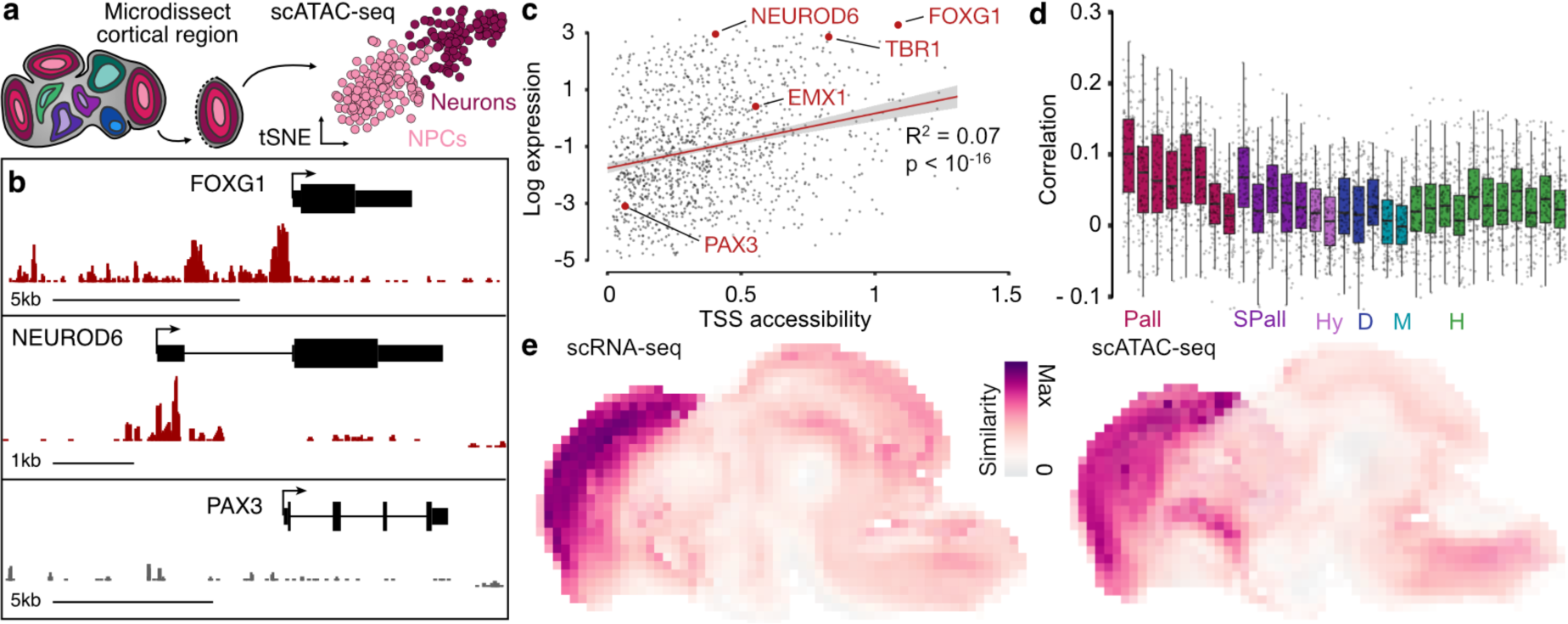
Projecting chromatin accessibility profiles to spatial brain maps. a) Single-cell RNA-seq (Camp et al., 2015; Kanton et al., 2019) and single-cell ATAC-seq (Kanton et al., 2019) was performed on single cortical regions microdissected from cerebral organoids between 2 and 4 months of age. b) Signal tracks showing accessibility (normalized read count) at the transcription start site (TSS) of forebrain marker FOXG1, cortex neuron marker NEUROD6, and hindbrain marker PAX3. c) Expression of all measured genes in the E13.5 pallium (cortex) as a function of TSS accessibility in cortical organoid cells. The fit of a linear model is indicated in red. d) Pearson correlation of organoid TSS accessibility (log normalized peak count) to gene expression in different brain structures. e) Sagittal projections of the E13.5 mouse brain colored by scaled similarity scores of organoid TSS accessibility (scATAC-seq) or expression (scRNA-seq) with voxel map expression of the downstream gene. Pall: pallium, SPall: subpallium, Hy: hypothalamus, D: diencephalon, M: midbrain: H: hindbrain.

Altogether, we have developed a set of computational tools that enables researchers to quickly and flexibly explore the vast amount of data within the Allen Brain Atlas resource. VoxHunt will be a powerful approach to assess novel organoid engineering protocols, to annotate cell fates that emerge in organoids during genetic and environmental perturbation experiments, and to develop data-driven hypotheses about the mechanisms underlying mammalian brain patterning that can be tested *in vitro*.

## METHODS

### Acquisition and preprocessing of Allen Brain Atlas in situ hybridization data

We downloaded 3D expression grid data from all *in situ* hybridization (ISH) experiments corresponding to the eight developmental time points (E11.5, E13.5, E15.5, E18.5, P4, P14, P28, P56) using the application programming interface (API) provided by the Allen Brain Atlas (ABA) Developing Mouse Brain Data Portal (http://help.brain-map.org/display/devmouse/API). In order to obtain in situ fluorescence intensity profiles (all genes) for each voxel, the data for each timepoint was summarized into a voxel by gene matrix. If multiple experiments were available for the same gene and time point, we calculated the arithmetic mean across experiments. The structure ontology was obtained by downloading the structure graph with ID 17 through the ABA web API (http://api.brain-map.org/). The resulting json file was further converted into a data frame. For differential expression testing and visualization, we introduced four additional levels of annotation by summarizing existing annotations. In these custom annotations, pallium (dorsal forebrain), subpallium (ventral forebrain), hypothalamus, diencephalon, midbrain, hindbrain and spinal cord are on the same level on the ontology tree and each splits further into finer structures. For the analysis shown here as well as for the VoxHunt package, we further removed all voxels annotated as spinal cord, as this structure was only included in the earliest developmental stages. The final expression matrices including metadata and custom annotations can be found as loom files under http://doi.org/10.17632/n6488nxzbh.1.

### Visualization of the structure annotation ontology

To visualize the structure ontology as a tree, we used the data.tree R package to convert the structure ontology data frame into a phylo object. We used the tidytree R package to manipulate the tree and add custom annotations to the nodes. Finally, the tree was plotted with the ggtree R package (Yu et al., 2017) using a circular layout and edges were colored according to our custom annotation level 3.

### Acquisition of organoid single-cell gene expression and chromatin accessibility data

With the goal of covering a range of different organoid protocols, we collected published single cell gene expression data from dorsal forebrain (Birey et al., 2017), ventral forebrain (Birey et al., 2017), thalamus (Xiang et al., 2019), and cerebral (Klaus et al., 2019) (unpatterned) organoids. Single cell gene expression and chromatin accessibility data for these organoids was obtained through the following GEO accessions: GSE93811 for Birey et al. (cortical and ventral), and GSE122342 for Xiang et al. (thalamus) and the following ArrayExpress accessions: E-MTAB-7552 for Kanton et al. (cerebral scRNA-seq) and E-MTAB-8089 for Kanton et al. (cerebral scATAC-seq).

### Single-cell gene expression data preprocessing

Transcript count matrices were preprocessed separately for each dataset with consistent parameters using the R package Seurat 3.0 (Stuart et al., 2019). Cells were filtered based on unique molecular identifier (UMI) counts (>500, <50000), the number of detected genes (>200, <6000) and the fraction of mitochondrial genes (<0.1). For all datasets, we annotated cells based on the cell type labels from the original paper and filtered out unannotated cells. Counts for each cell were normalized by the total number of counts for that cell, multiplied by a scaling factor of 10000 and subsequently natural-log transformed (NormalizeData()). Log normalized expression values were used as an input to the function FindVariableFeatures() to find the 2000 most variable genes based on the ‘vst’ selection method. Expression levels of these genes were used to perform Principal Component Analysis (PCA) with the function prcomp_irlba() from the irlba R package and the top 15 principal components (PCs) were used to produce a two-dimensional embedding with Uniform Manifold Approximation and Projection (UMAP) (Becht et al., 2018).

### Differential expression analysis

To perform differential expression analysis (DE) on both ISH and single-cell expression data, we calculated the Wilcoxon rank sum test p-values, AUROC values and log fold changes using the wilcoxauc() function from the R-package presto (Korsunsky et al., 2019). For the ISH data, we first performed DE on log2-transformed intensity values with a pseudocount of 1×10^−10^ between custom annotation levels and sorted by AUROC value to find highly specific marker genes for each structure (structure markers). To find time point-specific genes, we further performed DE between developmental stages for each structure on custom annotation level 2 separately (stage markers). Additionally, we searched for markers whose expression in a structure starts at any fetal developmental stage and is maintained in subsequent stages. For fetal stages E13.5, E15.5 and E18.5 we performed DE between the current stage and all and older stages versus all younger stages (e.g. for genes turned on at E15.5: E11.5/E13.5 vs E15.5/E18.5) for each structure on custom annotation level 2 separately (fetal development markers). For single-cell data, we performed DE on log normalized expression levels between annotated cell types for each dataset separately. Here, we determined unbiased cluster markers by taking the 3 genes with highest log fold change for each cluster.

### Correlation of single-cell transcriptomes to brain structures

To determine best matching brain structures for each neuronal cluster in the organoid datasets, we calculated the correlation of single cell transcriptomes with genes expression of structures in the fetal E13.5 mouse brain. For each cell in each cluster we calculated the Pearson correlation coefficient between the log normalized transcript count for each cell and the expression vector of each voxel in the E13.5 mouse brain based on 189 structure marker genes (top 10 for each structure at level 3). Correlation coefficients for each cell were then summarized for each structure by calculating the arithmetic mean. From the mean correlation coefficient *r*_*ij*_ of each cell *i* with each structure *j* we then defined the similarity to the structure as

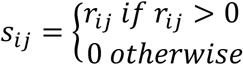

We assigned each cell to the structure with maximum similarity and displayed the mean similarity to each structure for all clusters.

### Selection of developmental stage

To determine the optimal developmental stage for spatial mapping, we calculated for each cell in each cluster the highest correlating voxel in the best matching structure of custom annotation level 2 for that cluster. We selected genes that were informative with regard to the developmental stage for each structure separately by taking the union of the above mentioned fetal development markers (top 50 AUROC for each stage and structure) and stage markers specific to E11.5 (top 50 AUROC for each structure) and used their expression values as an input for computing the Pearson correlation coefficient. In total, this resulted in 183 genes for the pallium, 181 genes for the subpallium, 179 genes for the midbrain and 181 genes for the diencephalon.

### Spatial correlation maps of organoid cells transcriptomes

To compute spatial correlation patterns of single-cell transcriptomes to the E13.5 mouse brain, we first removed background signal from *in situ* hybridization intensities by applying a minimum intensity threshold of 1. We further selected the 10 most specific markers for each structure at custom annotation level 3 to obtain a total of 201 genes. Based on the expression values of these genes, we calculated the Pearson correlation coefficient between each cell and each voxel in stage E13.5. Correlation coefficients for each cell cluster were summarized by computing the arithmetic mean for each voxel and subsequently transformed to a similarity metric as described above. Scaled correlation values were displayed as similarity maps across the brain in a sagittal view and on coronal sections. For direct cell-to-voxel projection, we further assigned each cell to the voxel with maximum correlation considering the 8 central sections (5-12) in the E13.5 mouse brain.

### Spatial correlation maps of organoid cells chromatin accessibility profiles

To compute correlation coefficients between transcriptome and chromatin accessibility profiles, the latter was summarized for each gene by counting all peaks in its TSS region (TSS ± 3kb) using the R package CHIPseeker (Yu et al., 2015). TSS peak counts were further normalized by the total number of counts for that cell, multiplied by a scaling factor of 10000 and subsequently natural-log transformed. We further selected the 10 most specific markers for each structure at custom annotation level 3 to obtain a total of 182 genes and used them to compute the Pearson correlation-based similarity between voxel expression values and log normalized gene accessibility scores for each cell as described above.

### Code availability

VoxHunt was implemented as an R package and is available at https://github.com/quadbiolab/voxhunt.

## SUPPLEMENTARY INFORMATION

**Figure S1:**
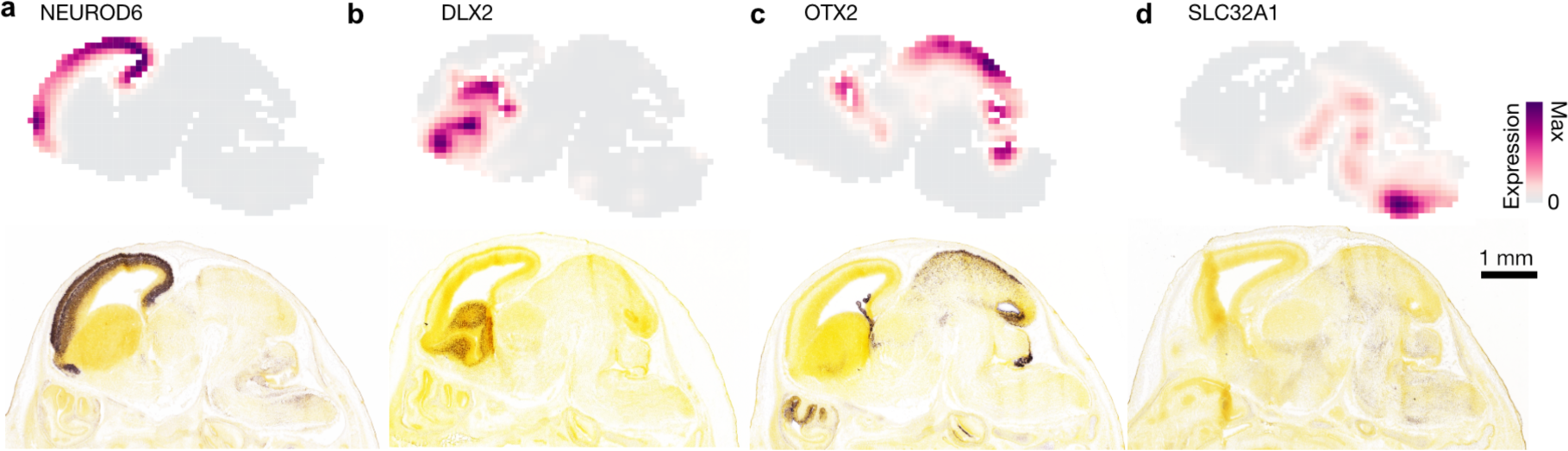
In silico expression maps compared to in situ hybridization patterns. Voxelized expression heatmaps (top) and *in situ* hybridization head images from the Allen Developing Mouse Brain Atlas (bottom) at E15.5 for DLX2 (a), NEUROD6 (b), OTX2 (c), and SLC32A1 (d).

**Figure S2:**
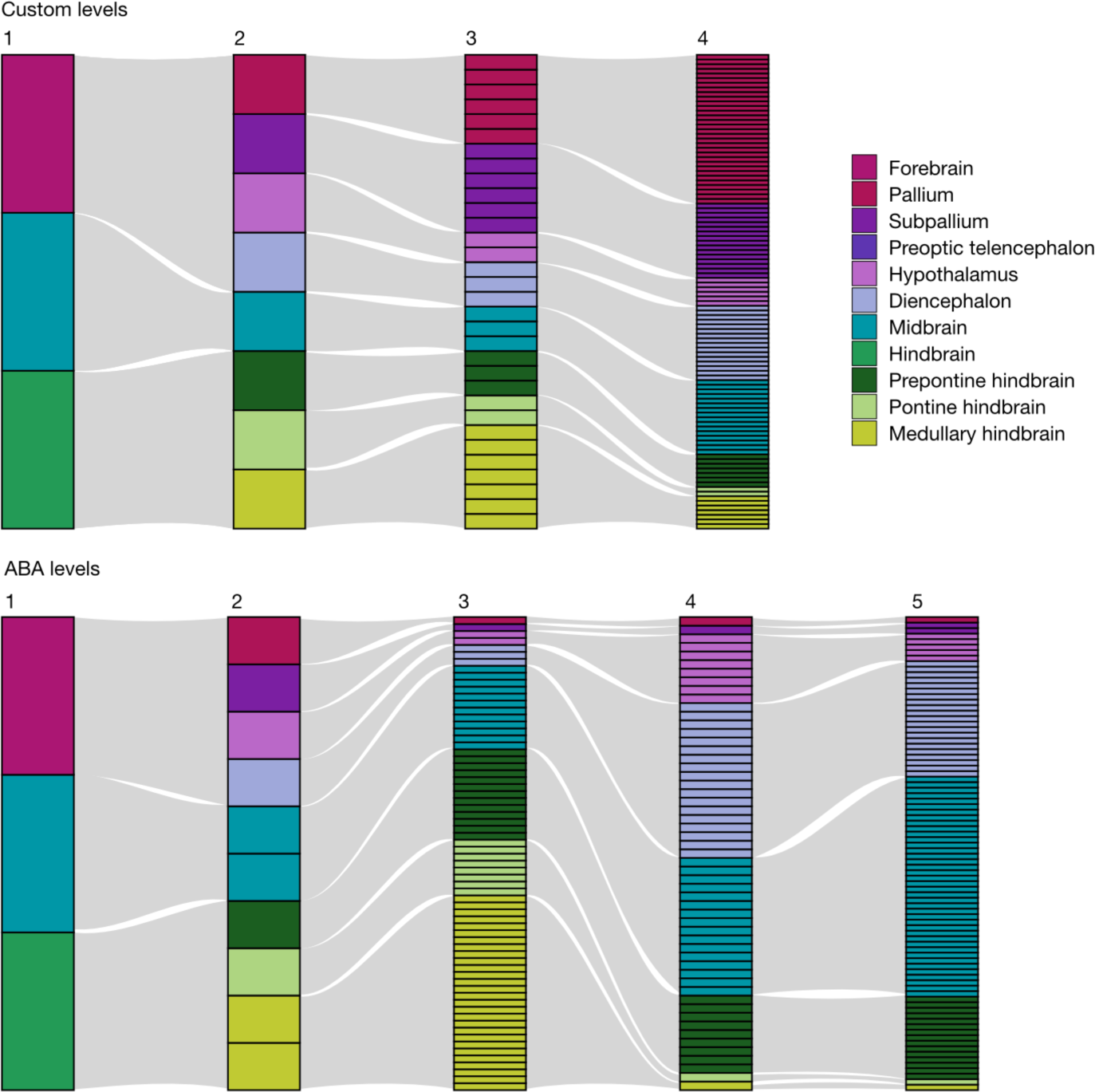
Hierarchical annotation levels of the developing mouse brain. Barcharts show how large structures are hierarchically subdivided into many smaller structures for custom annotation levels (top) and the first five annotation levels from the Allen Brain Atlas (bottom). Each box represents one annotated structure, colors represent custom level 1 on the first level and custom level 2 on all subsequent levels.

**Figure S3:**
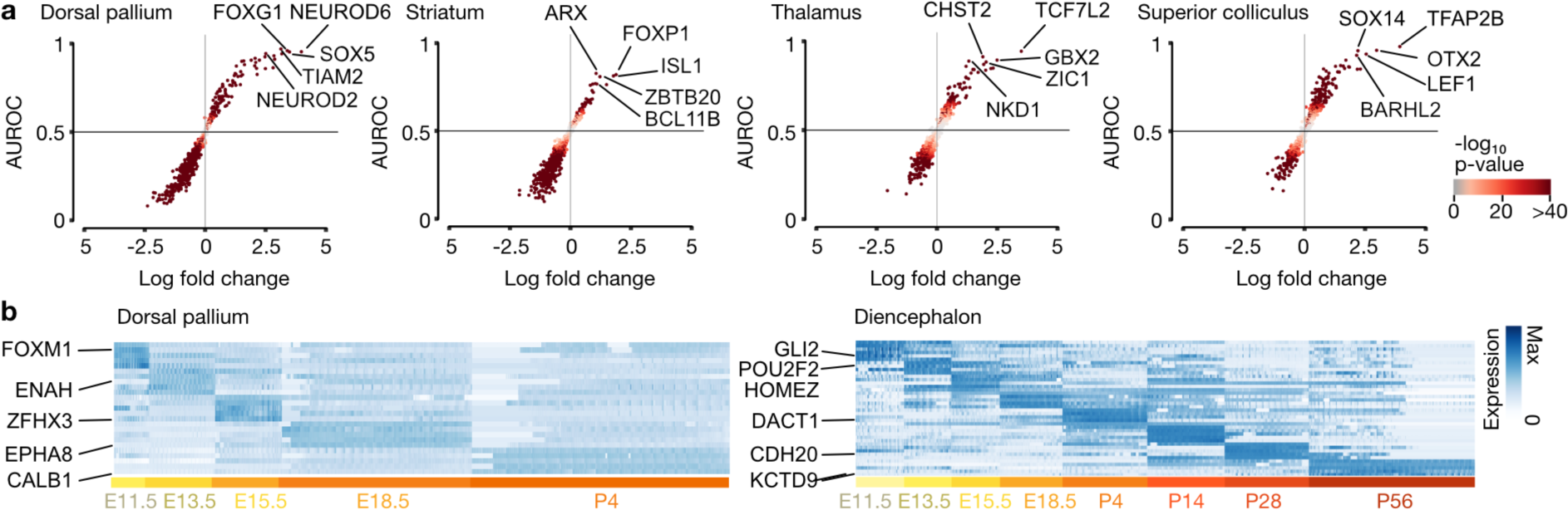
Feature extraction from voxel maps of developing mouse brain in situ hybridization data. a) Scatter plot showing feature selection for dorsal pallium, striatum, thalamus and superior colliculus. X-axis represents fold change and y-axis represents the area under the receiver operating characteristic curve (AUROC) metric for comparisons between the selected structure and all other brain structures. b) Heatmaps showing stage-specific genes for dorsal pallium and diencephalon binned by expression in the corresponding brain structure at different timepoints during development.

**Figure S4:**
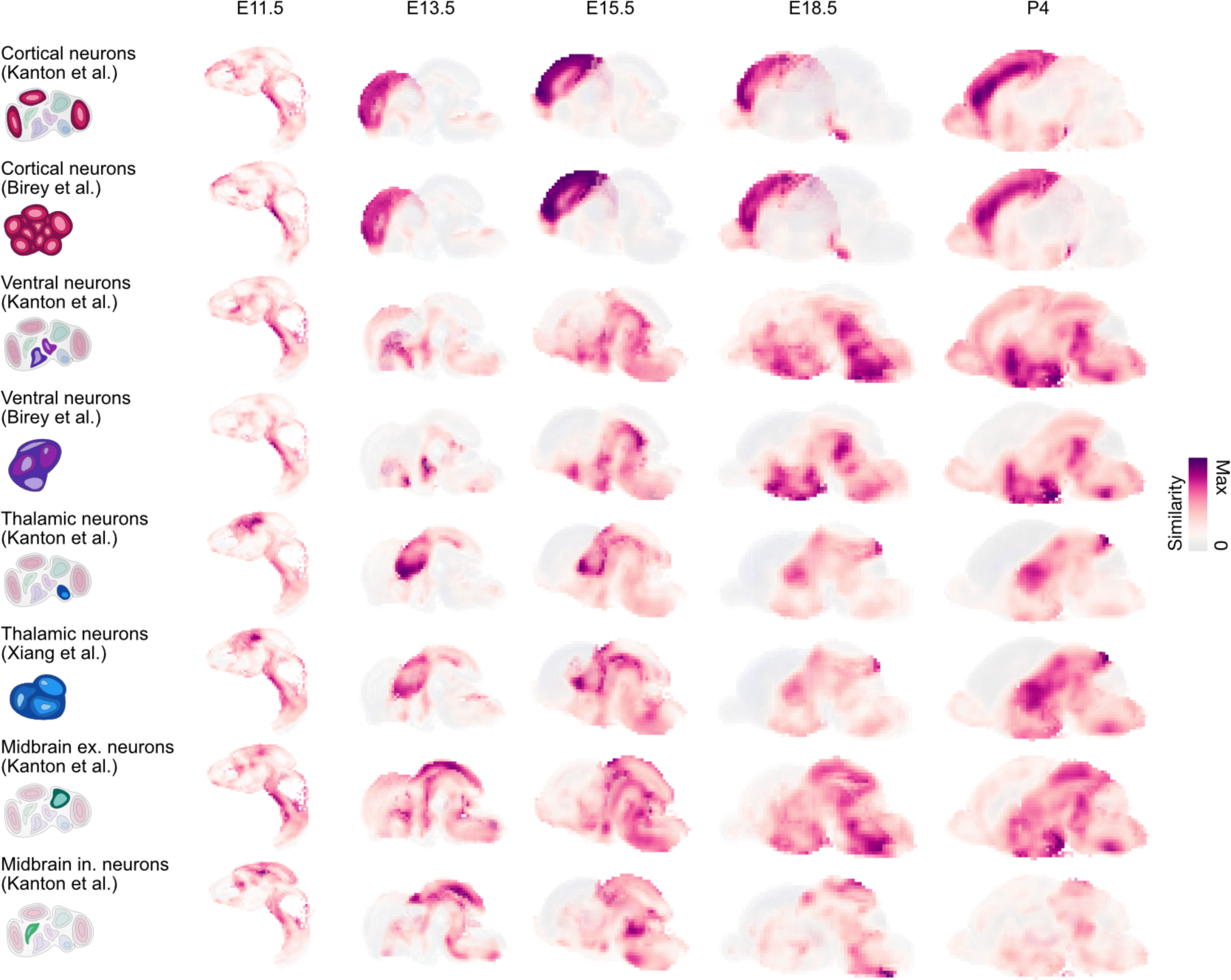
Spatial similarity maps of organoid cells across developmental stages of the developing mouse brain. Sagittal projections colored by scaled similarity scores of different organoid clusters from each organoid dataset to voxel maps of the developing mouse brain in stages E11.5 -P4.

**Figure S5:**
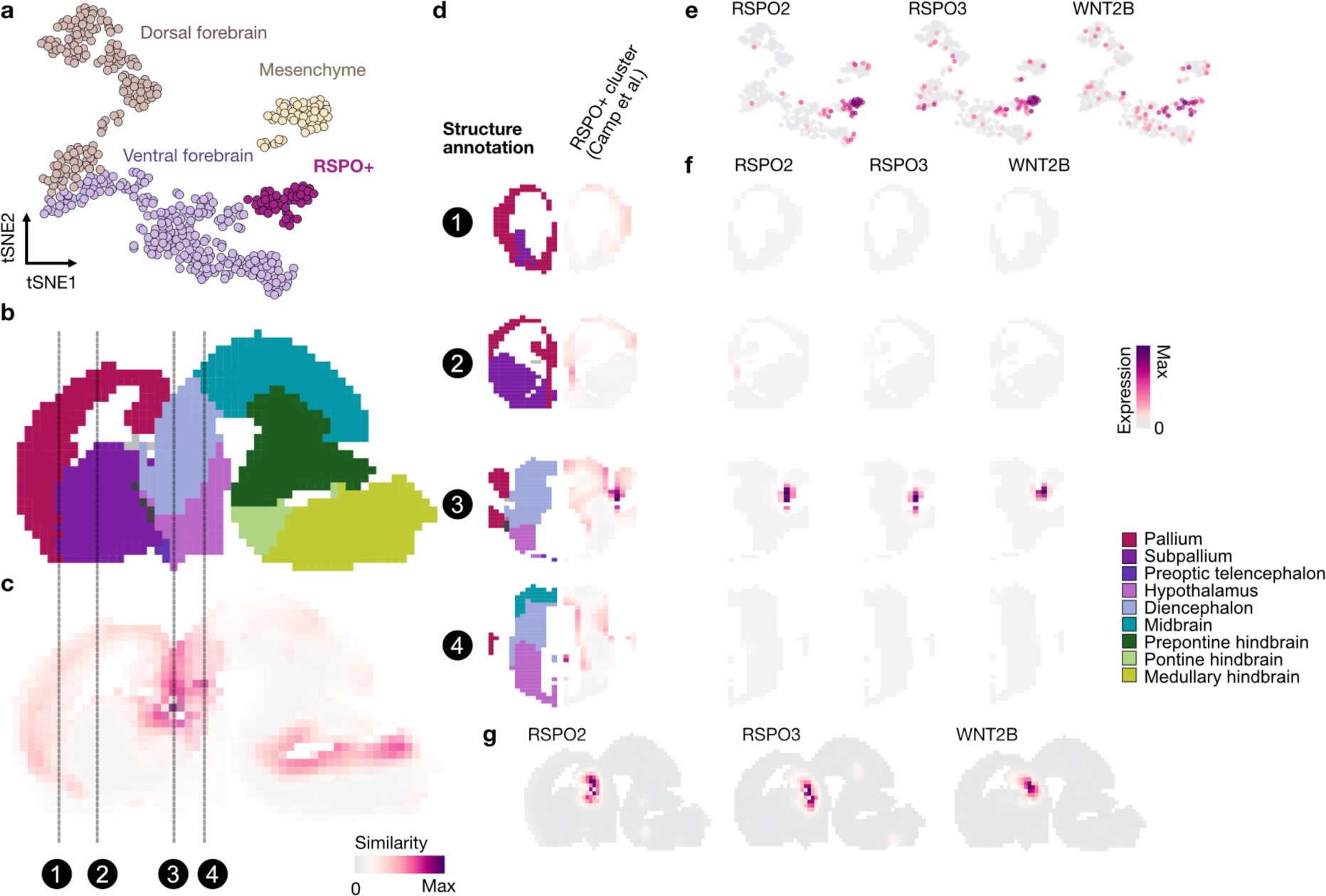
Spatial similarity maps of organoid cells reveal spatially confined cell types. a) tSNE embedding of single-cell transcriptomes from cerebral organoid cells based on the SMART-SEQ protocol implemented using Fluidigm C1 (Camp et al., 2015). The previously unknown ‘RSPO+’ cluster is highlighted in purple. b) Structure annotation of sagittal sections 2-5 of the E13.5 mouse brain. c) Sagittal projections colored by scaled similarity scores of the ‘RSPO+’ cluster to voxel maps of the E13.5 mouse brain. d) Coronal sections (6, 11, 22, 26) visualizing structure annotation and scaled similarity scores to different brain structures. e-f) Expression patterns of genes marking the ‘RSPO+’ cluster as maximum intensity projections across sagittal sections 2-5 (e) and coronal section 22 (f).

